# Acute systemic loss of Mad2 leads to intestinal atrophy in adult mice

**DOI:** 10.1101/2020.12.03.404186

**Authors:** Klaske M. Schukken, Yinan Zhu, Petra L. Bakker, Liesbeth Harkema, Sameh A. Youssef, Alain de Bruin, Floris Foijer

**Affiliations:** European Research Institute for the Biology of Ageing (ERIBA), University of Groningen, University Medical Center Groningen, 9713 AV, Groningen, the Netherlands; Dutch Molecular Pathology Center, Department of Biomolecular Health Sciences, Utrecht University, Faculty of Veterinary Medicine, 3584 CL, Utrecht, the Netherlands; Department of Pediatrics, University of Groningen, University Medical Center Groningen, 9713 AV, Groningen, the Netherlands

**Keywords:** Chromosomal instability, aneuploidy, CIN, cancer, mouse model

## Abstract

Chromosomal instability (CIN) is a hallmark of cancer, leading to aneuploid cells. To study the role that CIN plays in tumor evolution, several mouse models have been engineered over the last two decades. These studies have unequivocally shown that systemic high-grade CIN is embryonic lethal. We and others have previously shown that embryonic lethality can be circumvented by provoking CIN in a tissue-specific fashion. In this study, we provoke systemic high-grade CIN in adult mice as an alternative to circumvent embryonic lethality. For this, we disrupt the spindle assembly checkpoint (SAC) by alleviating Mad2 or truncating Mps1, both essential genes for SAC functioning, with or without p53 inactivation. We find that disruption of the SAC leads to rapid villous atrophy, atypia and apoptosis of intestinal epithelia, substantial weight loss, and death within 10 days after the start of the CIN insult. Despite this severe intestinal phenotype, most other tissues are unaffected, except for minor abnormalities in spleen, presumably due to the low proliferation rate in these tissues. We conclude that high-grade CIN *in vivo* in adult mice is most toxic to intestinal epithelia, presumably due to the high cell turnover in this tissue.

## Background

Chromosomal instability (CIN) is a process that leads to cells with an abnormal DNA content, a state also known as aneuploidy. The Spindle Assembly Checkpoint (SAC) helps to prevent CIN by arresting cells in metaphase until all chromosomes are properly attached to the spindle network. Inhibiting any protein involved in the SAC leads to high levels of CIN (1–3), including the kinase Mps1, which regulates checkpoint protein binding to kinetochores and Mad2, which inhibits the anaphase promoting complex (APC/c) when bound to kinetochores (4–10).

CIN is a hallmark of cancer cells (1,11–13), and while various mechanism can lead to chromosome miss-segregation (5,14–20), SAC alleviation is commonly used as a tool to provoke CIN in model systems. While SAC genes are rarely mutated (21), SAC function has been found to be impaired in various human cancers (14,21,22). Several mouse models have been engineered to study the effect of CIN on cancer initiation and progression (1,14). These models have unequivocally shown that systemic high-grade CIN is incompatible with embryonic development and further that low-grade CIN predisposes to cancer, most efficiently when combined with other predispositions such as p53 loss or APC mutation (1–3,23). To circumvent CIN-induced embryonic lethality (5,24), conditional mouse models were engineered in which CIN could be provoked in a tissue-specific fashion to study tolerance for chromosome mis-segregations in individual tissues *in vivo* (1,5).One intriguing finding from these models is that high levels of CIN are tolerated by some tissues, but not all. For instance, inducing CIN by inactivating Mad2 is incompatible with embryonic development (1), often lethal to cultured cells (25), and toxic to hair follicle stem cells (26), but tolerated by basal epidermal cells (26), T-cells (27), and hepatocytes *in vivo* (27). Furthermore, systemic CIN provoked in adult mice drives tumorigenesis in a dose-dependent fashion with medium CIN rates being most efficient (23). Here, we study the effects of acute high-grade CIN in adult mice provoked by complete SAC alleviation and find that this leads to rapid regression of intestinal epithelia followed by significant weight loss and ultimately death.

## Results

CIN driven by systemic Mad2 inactivation is incompatible with mouse embryonic development (24). However, the impact of acute systemic Mad2 loss in adult mice, circumventing embryonic defects, has not been tested. For this purpose, we crossed *Mad2*^*f/f*^ (26,27) and *Mps1*^*f/f*^ (10) conditional mice with mice expressing a ubiquitous tamoxifen-inducible Cre recombinase (*Cre-ERT2*, (33)). In these mice, tamoxifen treatment yields acute and systemic Mad2 loss. Since p53 loss can partly rescue Mad2-loss imposed cell death *in vivo* and in cultured ES cells (6), we also crossed the Mps1^f/f^;Cre-ERT2 and Mad2^f/f^;Cre-ERT2 strains into a p53 conditional knockout background.

Next, we compared protocols for tamoxifen administration: intraperitoneal (IP) injections, oral gavage and tamoxifen administration through food pellets and compared the effects of the Mps1 and Mad2 alleles as CIN drivers, with and without p53 conditional alleles (**Sup. Fig 1**). We did not observe significant differences between the administration route of tamoxifen, the CIN-driving alleles, nor p53 status: most mice had to be sacrificed because of excessive weight loss within two weeks (**Sup. Fig 1**). As IP injections come with a risk of intestinal punctures and mouse food intake was reduced for tamoxifen-containing food pellets, also in control mice, in follow up experiments, we administrated tamoxifen by oral gavage. Furthermore, as we did not observe any differences in the phenotypes of mice with *Mad2*^*f/f*^ and *Mps1*^*f/f*^ alleles, we only pursued Mad2 conditional mice.

To understand the cause of the excessive weight loss, we next setup a cohort of 4 *Mad2*^*f/f*^ Cre-ERT2 and 4 *Mad2*^*f/f*^;*p53*^*f/f*^;*Cre-ERT2* mice. As controls, we included two Cre‐ERT2 mice that received control-vehicle instead of tamoxifen, and two Cre‐ERT2 negative mice that received tamoxifen. While control mice gained an average of 7% of their body weight; *Mad2*^*f/f*^;*Cre‐ERT2* and *Mad2*^*f/f*^;*p53*^*f/f*^;*Cre‐ERT2* mice lost 14 and 13% of their weight, respectively within four days after treatment had started (**Fig. 1A**).

**Figure 1:**
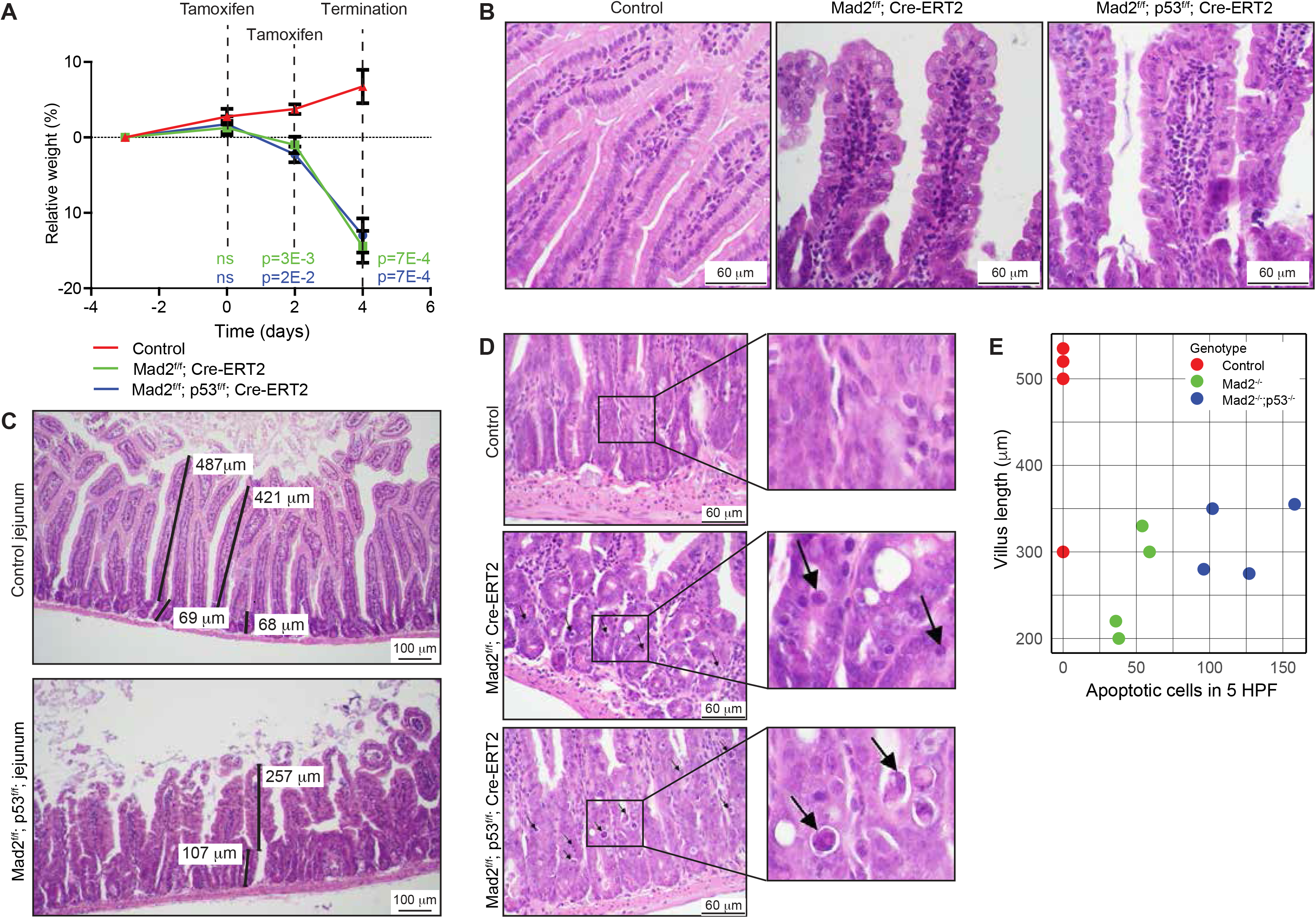
Inactivation of Mad2 knockout leads to rapid atrophy of intestinal epithelia. (**A**) Relative weight compared to starting weight over time. Plotted average weight with SEM, n=4 per condition. Two-sided t‐ test was used to test difference between each genetic group and the control, p‐value per timepoint and per genotype displayed. (**B**) Representative images of jejunum of control and Mad2; p53 compound knockout mice. H&E staining, scale bar is 60 μm. (**C**) Representative images of villi and crypts of the jejunum. H&E staining, scale bar = 100 μm. i) normal jejunum ii) jejunum with atrophied villi and atypical cells. (**D**) Representative images of villi and crypts of the jejunum per genotype. H&E staining, scale bar is 60 μm. Black arrows point to apoptotic cells. (**E**) Scatter plot of villus length and apoptotic cells per High Power Field (HPF) image per genotype.

Next, mice were euthanized and tissues harvested for analysis. We inspected the intestinal jejunum, spleen, lung, liver, kidneys, and a blood smear for each mouse (**Table 1, 2**). When evaluating the intestinal epithelia, we noticed several atypia of the mucosal cells of the villi, including abnormal columnar shape and karyomegaly and/or binucleated cells in the knockout mice, the latter fitting well with chromosomal instability imposed by Mad2 loss (**Fig. 1B**). Most strikingly, the knockout mice displayed a significant decrease in villus length in the intestinal jejunum (**Fig. 1C**), which coincided with significant increase in apoptotic cells (**Fig. 1D, Table 1**). While survival or weight loss kinetics between p53-proficient and - deficient mice were comparable, *Mad2*^*f/f*^;*p53*^*f/f*^ *Cre-ERT2* mice displayed a higher number of large apoptotic cells with an abnormal shape and less condensed chromatin in jejunum than *Mad2*^*f/f*^ mice, suggesting an increased frequency of mitotic catastrophe in *Mad2*^*f/f*^;*p53*^*f/f*^;*Cre-ERT2* mice (**Fig. 1D, E**), a p53-independent type of apoptosis (34). The observed phenotype is consistent with the high turnover of intestinal cells: intestinal villi proliferate rapidly and are renewed every 3‐5 days (35). The atrophy and cell death most likely interfered with the uptake of nutrients, causing extreme weight loss ultimately leading to death of the mice.

**Table 1:** Histological data for intestine per genotype **Table 1 Legend**: **Villus atrophy** (44): Scores: 0= absent; 1= mild: ~75% of normal length; 2= moderate: ~50% of normal length; 3= Marked: <25% of normal length **Villus epithelium degeneration** Scores: 0= absent; 1= mild: Epithelial attenuation, loss of brush border; 2= moderate: clear epithelial degeneration; 3= severe: Loss of epithelial cells **Villus epithelium atypia** Scores: 0= absent, 1= mild, 2= moderate, 3= severe **Crypt epithelial degeneration** Scores: 0= absent; 1= mild: Epithelial attenuation; 2= moderate: clear epithelial degeneration; 3= severe: Sloughing and loss of epithelial cells **Crypts**(44): Scores: 0= normal; 1= 10%; 2= 10-50%; 3= >50% of the crypts contain apoptotic cells/debris

| Genotype | Mouse ID | Intestine |  |  |  |  |  |  |  | Apoptotic cells in 5 HPF |
| --- | --- | --- | --- | --- | --- | --- | --- | --- | --- | --- |
|  |  | Crypt depth (um) | Villus length (um) | Villus atrophy | Villus epithelium degeneration | Villus epithelium atypia | Crypt epithelial degeneration | Crypt epithelial atypia | Crypts |  |
| Mad2<sup>f/f</sup> p53<sup>f/f</sup> | 118 | 100 | 350 | 1 | 1 | 1.5 | 2 | 2.5 | 3 | 102 |
| Mad2<sup>f/f</sup> p53<sup>f/f</sup> | 121 | 115 | 275 | 2 | 1 | 2.5 | 2 | 2.5 | 3 | 127 |
| Mad2<sup>f/f</sup> p53<sup>f/f</sup> | 123 | 85 | 355 | 1 | 1 | 1.5 | 2 | 2 | 3 | 158 |
| Mad2<sup>f/f</sup> p53<sup>f/f</sup> | 135 | 90 | 280 | 2 | 2 | 3 | 2.5 | 3 | 3 | 96 |
| Mad2<sup>f/f</sup> | 127 | 90 | 330 | 1 | 1.5 | 2.5 | 2 | 3 | 2 | 54 |
| Mad2<sup>f/f</sup> | 129 | 80 | 220 | 2.5 | 0 | 3 | 3 | 2.5 | 3 | 36 |
| Mad2<sup>f/f</sup> | 131 | 100 | 300 | 1 | 1 | 1 | 2 | 2 | 2 | 59 |
| Mad2<sup>f/f</sup> | 143 | 80 | 200 | 2.5 | 1 | 3 | 2.5 | 3 | 3 | 38 |
| control | 138 | 105 | 520 | 0 | 0 | 0 | 0 | 0 | 0 | 0 |
| control | 139 | 85 | 300 | 1 | 0 | 0 | 0 | 0 | 0 | 0 |
| control | 140 | 105 | 535 | 0 | 0 | 0 | 0 | 0 | 0 | 0 |
| control | 125 | 80 | 500 | 0 | 0 | 0 | 0 | 0 | 0 | 0 |
|  |  | **t.test** |  | **Wilcox sum rank test** |  |  |  |  |  |  |
| Mad2<sup>f/f</sup> p53<sup>f/f</sup> vs. control |  | 0.70 | 0.046 | 0.047 | 0.018 | 0.020 | 0.018 | 0.020 | 0.013 | 0.021 |
| Mad2<sup>f/f</sup> vs. control |  | 0.47 | 0.019 | 0.047 | 0.067 | 0.020 | 0.020 | 0.020 | 0.019 | 0.021 |

**Table 2:** Histological data of organs per genotype: Spleen, Blood. Lung Liver and Kidneys. **Table 2 Legend**: **Spleen Atypical: Atypical cells within the red pulp** Scores: 0= absent; 0.5= rarely single atypical cell present; 1= 1-5 per HPF; 2= 5-10 per HPF; 3=10-20 per HPF **Spleen EMH (extramedullary hematopoiesis within the red pulp)** Scores: 0= absent, 1= mild, 2= moderate, 3= marked **Abbreviations** nsl= No Significant Lesions HPF= High Power Fields. The area visible under a high magnification microscope at 400X.

| Genotype | Mouse ID | Spleen |  |  | Other tissues |  |  | Blood |  |  |  |
| --- | --- | --- | --- | --- | --- | --- | --- | --- | --- | --- | --- |
|  |  | Spleen Atypical | Spleen EMH | Spleen Follicles | Lung | Liver | Kidney | Rbc diameter (µm) | Polychromatophilia | Size of polychromatic cells (µm) | Howell-Jolly bodies |
| Mad2<sup>f/f</sup> p53<sup>f/f</sup> | 118 | 3 | 1 | nsl | nsl | nsl | nsl | 6-6.5 | 0 |  | 4 / HPF |
| Mad2<sup>f/f</sup> p53<sup>f/f</sup> | 121 | 3 | 1 | nsl | nsl | nsl | nsl | 6-6.5 | 1 / HPF |  | 4 / HPF |
| Mad2<sup>f/f</sup> p53<sup>f/f</sup> | 123 | 2.5 | 1.5 | nsl | nsl | nsl | nsl | 5.5-6 | 0 |  | 1 / HPF |
| Mad2<sup>f/f</sup> p53<sup>f/f</sup> | 135 | 2 | 1 | nsl | nsl | nsl | nsl | 6 | 0 |  | 3 / HPF |
| Mad2<sup>f/f</sup> | 127 | 0.5 | 1.5 | nsl | nsl | nsl | nsl | 5.5-6 | 0 |  | 3 / HPF |
| Mad2<sup>f/f</sup> | 129 | 0 | 1.5 | nsl | nsl | nsl | nsl | 6-6.5 | 0 |  | 2 / HPF |
| Mad2<sup>f/f</sup> | 131 | 0.5 | 1 | nsl | nsl | nsl | nsl | 5.5-6 | 0 |  | 1 / HPF |
| Mad2<sup>f/f</sup> | 143 | 0 | 0 | nsl | nsl | nsl | nsl | 6-6.5 | 0 |  | 4 / HPF |
| control | 138 | 0 | 2 | nsl | nsl | nsl | nsl | 6-6.5 | 6 / HPF | 7 | 1 / HPF |
| control | 139 | 0 | 2 | nsl | nsl | nsl | nsl | 6.5 | 20 / HPF | 6.5-7 | 6 / HPF |
| control | 140 | 0 | 2 | nsl | nsl | nsl | nsl | 5.5-6.5 | 25 / HPF | 6.5-7 | 2 / HPF |
| control | 125 | 0 | 2 | nsl | nsl | nsl | nsl | 6 | 13 / HPF | 6.5-7 | 1 / HPF |
|  |  | **Wilcox sum rank test** |  |  |  |  |  | **t.test** |  |  | **t.test** |
| Mad2<sup>f/f</sup> p53<sup>f/f</sup> *vs.* control |  | 0.020 | 0.018 |  |  |  |  | ns |  |  | ns |
| Mad2<sup>f/f</sup> *vs.* control |  | 0.018 | 0.020 |  |  |  |  | ns |  |  | ns |

In addition to the intestinal abnormalities, we also observed large, atypical cells within the red pulp of the spleen in *Mad2*^*f/f*^;*p53*^*f/f*^;*Cre-ERT2* mice (**Fig. 2A, B, Table 2**), and to a lesser extent in two of the four *Mad2*^*f/f*^;*Cre-ERT2* mice. These atypical cells showed a high nuclear/cytoplasm ratio with nuclear atypia and occasional binucleation, again in line with a CIN phenotype. While this is unlikely to be the cause of the extreme weight loss and death, these changes might thus represent mild hematopoietic dysplasia related to the first effects of Mad2 (and p53) loss in the hematopoietic system. We did not find abnormal cells in the blood smears, suggesting that the lymphoid cells in peripheral blood were not affected within the first 4 days of tamoxifen treatment. Finally, we did not observe any abnormalities in liver, lung and kidneys (**Table 2**), likely due to the much lower turnover of cells in these tissues and the short timeframe of the experiment.

**Figure 2:**
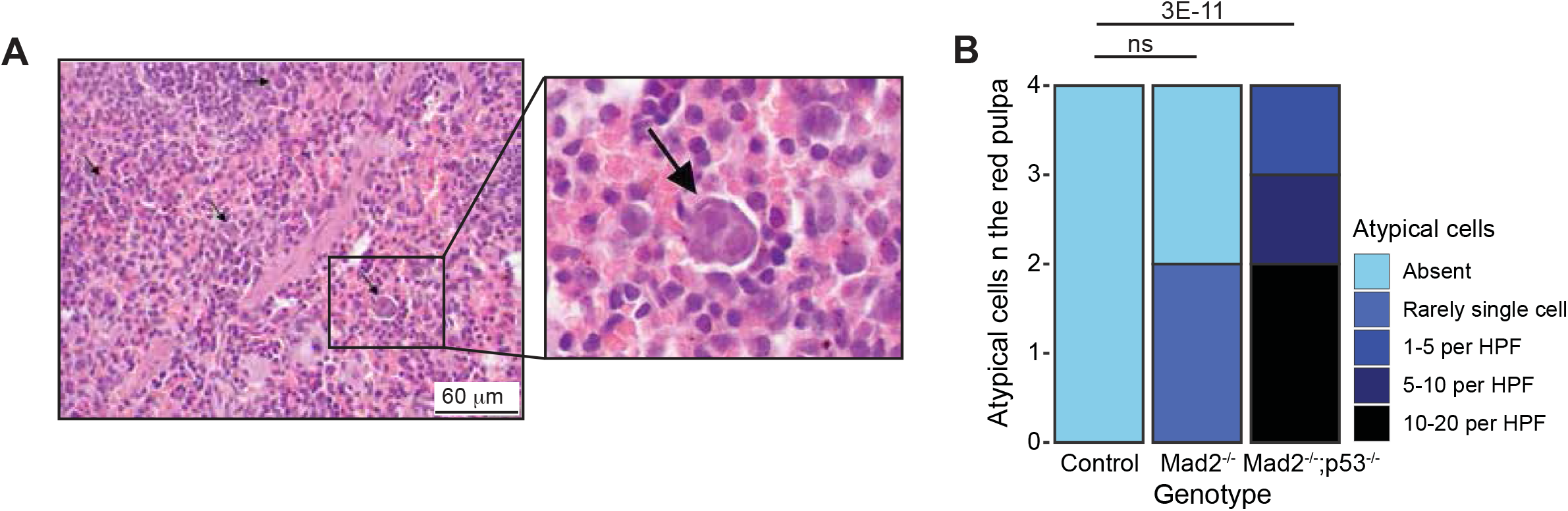
Acute Mad2 loss leads to a modest phenotype in spleen within the first four days (**A**) *Mad2*^*−/−*^ *p53*^*−/−*^ spleen showing cellular atypia in the red pulp, scale bar 20 μm, HE staining, magnification 400x. (B) Quantification of spleen abnormalities between genotypes, n=4 per genotype. p-values refer to Wilcox rank‐ sum test.

We conclude that acute systemic loss of Mad2 with or without p53 inactivation causes rapid atrophy of intestinal epithelia yielding reduced nutrient uptake, ultimately leading to death. While this phenotype is intriguing and suggests that Mad2 loss is particularly toxic to the stem cells residing in the intestinal crypts, similar to what was observed previously for hair follicle stem cells (26), the severity of the phenotype also precluded assessment of Mad2 loss in other adult tissues.

## Discussion

Systemic inactivation of the SAC, and the resulting high levels of CIN, lead to early embryonic death (1–3). However, to our knowledge, the consequences of complete systemic SAC alleviation in adult mice have not been reported so far. We have previously shown that tissue-specific inactivation of the SAC can circumvent embryonic lethality associated with Mad2 loss (26,27), and found that different tissues cope differently with SAC loss. For instance, inactivation in the epidermis revealed that SAC alleviation is not tolerated by hair follicle stem cells, but remarkably wel-tolerated by the basal cells of the epidermis (26). In this study, we find that systemic inactivation of the SAC leads to rapid death of adult mice coinciding with rapid weight loss. The rapid weight loss following Mad2 loss coincides with increased apoptosis of proliferating cells in the intestinal crypts and transient amplifying cells, leading to a severe atrophy of intestinal epithelia.

While we observed a significant phenotype in intestine, we did not find any noticeable effects of SAC loss in lung, liver, and kidneys within the four-day timeframe. We did observe a weak phenotype in the hematopoietic system, mostly in spleen. A possible explanation for this is the proliferation rate within these tissues, as both intestinal crypt cells and hematopoietic stem cells produce rapidly proliferating cells (35–37). Notably, while loss of p53, or p53 mutation enhances CIN tolerance in MEFs (6), we did not observe striking differences between the phenotypes of *Mad2*^*f/f-*^;*Cre-ERT2* and *Mad2*^*f/f*^;*p53*^*3f/f*^;*Cre-ERT2* adult mice, suggesting that in intestine p53 loss is not sufficient to rescue apoptotic cell death in a high-grade CIN background. Overall, our data shows that acute high-grade CIN is particularly toxic to tissues that exhibit a high cell turnover.

Since CIN is a hallmark of cancer cells (13,14), and three out of four tumors are aneuploid (11,12), it is of the utmost importance to know to what extent individual tissues tolerate CIN *in vivo*, as this will contribute to our understanding of cancer progression per tissue type. Our findings also have clinical implications as exacerbating CIN may be a very effective way to kill cells with a CIN phenotype (4,8,9,29,30,38–40), which is being investigated as a potential cancer therapy (28,41,42). Our finding that provocation of high-grade CIN trough SAC alleviation is very toxic to proliferative tissues should therefore be taken into account when further developing such strategies.

## Material and Methods

### Animal experiments

All animal experiments were performed in accordance with Dutch law and approved by the University of Groningen Medical Center Committees on Animal Care. The conditional *Mad2*^*f/f*^ and *Mps1*^*f/f*^ animal models were previously described (10,26,27). *Trp53*^*f/f*^ (*p53*^*f/f*^) conditional knockout mice were a gift from Jos Jonkers (43). All mice used were between 9-10 weeks old. *Mad2*^*f/f*^ or *Mps1*^*f/f*^ mice were bred with ubiquitous Cre‐ERT2 mice (33) and p53^f/f^ mice to obtain the various cohorts of mice described in this study.

### Tamoxifen inductions

To find the optimal method of tamoxifen administration, three different administration methods were compared in a small cohort of mice, which included tamoxifen provided in food pellets (Envigo, 250mg/kg), tamoxifen in peanut oil injected intraperitoneally or by oral gavage. As mice put on a tamoxifen-containing diet had lower food intake than mice on a normal diet, and IP injections come with a risk of intestinal punctures, we used oral gavage for the experiment cohort. For IP injections and oral gavage, we administrated 0.13 mg tamoxifen (Sigma) per gram of mouse in peanut oil (Sigma).

### Histological analysis

Animals were euthanized, organs removed and washed with PBS. Tissues were placed in a cassette, and fixed in 10% neutral buffered formalin for 1‐2 days, then moved into 70% ethanol. Histology slides were prepared and standard H&E staining was performed at the Department of Pathology at Utrecht University.

### Statistical analysis and Plots

Intestinal villi length, crypt length, red blood cell diameter and number of Howell‐Jolly bodies present in blood samples were compared with a two‐sided t‐test. A Wilcox rank sum test was used for categorical data, and data with values near zero, as indicated in Table 1 and 2. A two‐sided t‐test was used to evaluate the difference in mouse weight change (relative to day 0) per group, comparing experimental to control mice. A log‐rank (Mantel‐Cox) test was used to compare survival curves. Survival curves and weight loss graph were plotted in GraphPad PRISM..

## Acknowledgements

We would like to thank the members of the Foijer lab and Bruggeman lab for fruitful discussion.

---

**Supplementary Figure 1**: Kaplan-Meier curves per genotype for several tamoxifen administration routes. (A, B) Kaplan-Meier curves for (**A**) *Mad2*^*f/f*^ *p53*^*f/f*^ *Cre-ERT2*, or (**B**) *Mps1*^*f/f*^ *p53*^*f/f*^ *Cre-ERT2* and *Mps1*^*f/f*^ *Cre-ERT2* mice for various routes of tamoxifen administration. Number of mice per genotype/treatment type is listed.

