## Supplementary figures and images for "Acute systemic loss of Mad2 leads to intestinal atrophy in adult mice"

### Supplementary Figure 1

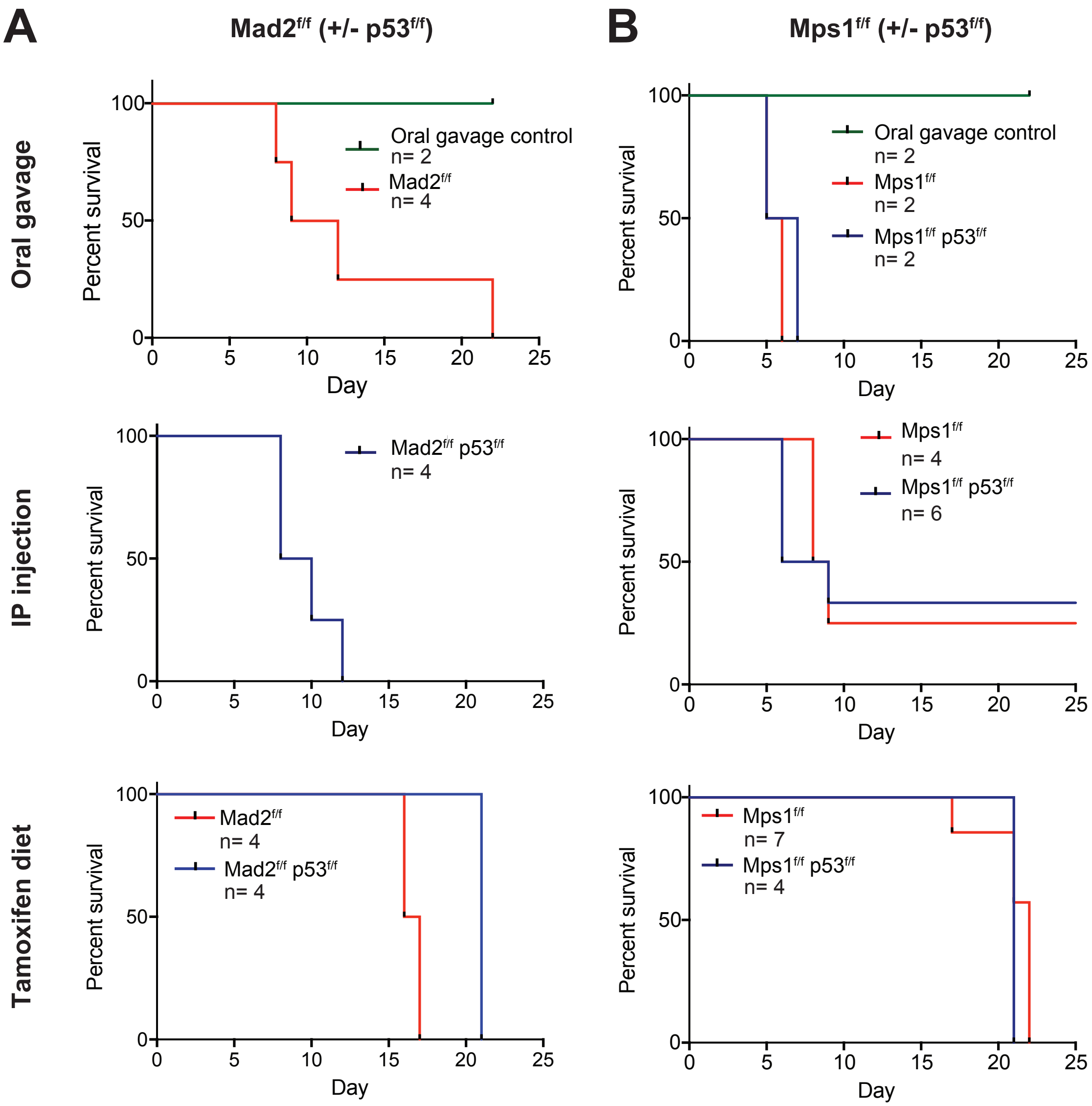
